# Discriminative aversive learning and amygdala responsivity is enhanced in mice with reduced serotonin transporter activity

**DOI:** 10.1101/154690

**Authors:** João Lima, Trevor Sharp, Amy M. Taylor, David M. Bannerman, Stephen B. McHugh

## Abstract

The serotonin (5-HT) transporter (5-HTT) regulates 5-HT availability at the synapse. Low or null 5-HTT expression results in increased 5-HT availability and has been reported to produce anxious and depressive phenotypes, although this remains highly controversial despite two decades of investigation. Paradoxically, SSRIs, which also increase 5-HT availability, reduce the symptoms of anxiety and depression. An emerging ‘network plasticity’ theory of 5-HT function argues that, rather than influencing mood directly, increasing 5-HT availability enhances learning about emotionally-significant events but evidence supporting this theory is inconclusive. Here, we tested one key prediction of this theory: that increased 5-HT availability enhances aversive learning. In experiment 1, we trained 5-HTT knock-out mice (5-HTTKO), which have increased 5-HT availability, and wild-type mice (WT) on an aversive discrimination learning task in which one auditory cue was paired with an aversive outcome whereas a second auditory cue was not. Simultaneously we recorded neuronal and hemodynamic responses from the amygdala, a brain region necessary for aversive learning. 5-HTTKO mice exhibited superior discrimination learning than WTs, and had stronger theta-frequency neuronal oscillations and larger amygdala hemodynamic responses to the aversive cues, which predicted the extent of learning. In experiment 2, we found that acute SSRI treatment (in naïve non-transgenic mice), given specifically before fear learning sessions, enhanced subsequent fear memory recall. Collectively, our data demonstrate that reducing 5-HTT activity (and thereby increasing 5-HT availability) enhances amygdala responsivity to aversive events and facilitates learning for emotionally-relevant cues. Our findings support the network plasticity theory of 5-HT function.

## Introduction

The serotonin (5-HT) transporter (5-HTT) regulates 5-HT availability at the synapse, and variation in 5-HTT expression is thought to influence emotional phenotypes. Genetic polymorphisms in the promoter region of the human 5-HTT gene produce variation in 5-HTT expression, with lower expression in short allele carriers (SS or SL) compared to long (LL) allele homozygotes, specifically the L_A_L_A_ subtype (Hu *et al*, 2006; Murphy *et al*, 2013). Compared to L_A_L_A_, short allele carriers have been reported to exhibit increased neuroticism, more reactive amygdala responses, and are at a higher risk of developing affective disorders, especially when combined with aversive life experiences (Caspi *et al*, 2010; Caspi *et al*, 2003; Hariri *et al*, 2005; Hariri *et al*, 2002; Lesch *et al*, 1996). Similarly, experiments in monkeys and genetically-modified rodents report that low (or null) 5-HTT expression results in higher 5-HT availability and anxious and fearful phenotypes (Bennett *et al*, 2002; Champoux *et al*, 2002; Holmes *et al*, 2003; Jennings *et al*, 2010; Mathews *et al*, 2004; Olivier *et al*, 2008; Wellman *et al*, 2007).

However, several large-sample human studies have found little or no influence of 5-HTT genotype on emotional phenotypes (Culverhouse *et al*, 2017; Middeldorp *et al*, 2010; Surtees *et al*, 2006; Willis-Owen *et al*, 2005). Furthermore, chronic treatment with SSRIs, which also increases 5-HT availability, paradoxically reduces the symptoms of anxiety and depression in humans and animal models. So does 5-HTT expression affect emotionality? And if so, what are the cognitive and neurobiological mechanisms that underpin any such relationship? Despite a substantial research effort, there is still no satisfactory answer to these questions. One potential resolution has come from the proposal that 5-HT availability does not regulate emotional state directly but rather it regulates plasticity in the neural networks underlying emotional learning and therefore influences sensitivity to emotional life events (Castren, 2005). Thus emotional experiences, both positive and negative, may be encoded more strongly in low 5-HTT-expressing individuals because they have higher 5-HT availability (Belsky *et al*, 2009; Branchi, 2011; Homberg and van den Hove, 2012).

One important prediction of the network plasticity theory is that reduced (or null) 5-HTT expression will facilitate learning of aversive experiences. Although some evidence is consistent with this prediction (Barkus *et al*, 2014; Garpenstrand *et al*, 2001; Line *et al*, 2014; Lonsdorf *et al*, 2009), there are notable exceptions. For example, 5-HTT knock-out (5-HTTKO) mice and rats reportedly exhibit essentially normal fear learning (Muller *et al*, 2011; Narayanan *et al*, 2011; Shan *et al*, 2014; Wellman *et al*, 2007). However, these studies used ‘single-cue’ conditioning paradigms (e.g. several exposures to an auditory cue paired with foot-shock), and wild-type animals typically displayed high levels of fear-related behavior (i.e. ‘freezing’), leaving only a small window to detect group differences. In contrast, human studies reporting an effect of 5-HTTLPR in fear learning have used discriminative paradigms which involve learning which of two stimuli are associated with an aversive outcome (Garpenstrand *et al*, 2001; Lonsdorf *et al*, 2009).

Here, we investigated whether 5-HTT expression in mice influenced aversive learning using a discriminative fear learning task. We trained 5-HTTKO mice and their wild-type littermates to discriminate between one auditory cue that was paired with foot-shock (aversive cue) and a second auditory cue that was never paired with foot-shock (non-aversive cue). Simultaneously, we recorded local field potentials (LFPs) and hemodynamic responses from the amygdala during fear conditioning to determine the effects of reduced 5-HTT activity on neural correlates of aversive learning (Barkus *et al*, 2014; McHugh *et al*, 2014; McHugh *et al*, 2013). In a second experiment, we tested whether transiently increasing 5-HT availability with selective serotonin reuptake inhibitors (SSRIs), specifically during conditioning, influenced fear learning and memory. We predicted that both 5-HTTKO and SSRI-treated mice would exhibit enhanced aversive learning.

## Materials and Methods

Detailed methods can be found in the supplementary material

### Subjects

Experiment 1 used 31 wild-type (WT) mice (female: n=16) and 29 5-HTTKO mice (female: n=14). Mice were generated as described previously (Bengel *et al*, 1998) and backcrossed onto a C57BL/6J background for at least eight generations. Experiment 2 used 96 female C57BL/6J mice (Charles River Laboratories, Kent, UK). Mice were 4-10 months (5-HTTKO and WT) or ~2 months old (C57BL/6J) at the start of testing. All experiments were conducted in accordance with the United Kingdom Animals Scientific Procedures Act (1986) under project licenses PPL 30/2561 and 30/3068, and were approved by local ethical review for the University of Oxford.

### Surgery

Under isoflurane anesthesia, mice (WT: n=14, female: n=8; 5-HTTKO: n=14, female: n=7) were implanted with electrodes into the basolateral amygdala (BLA) to record tissue oxygen (TO_2_) and local field potentials (LFPs). Co-ordinates were -1.35mm anterior/posterior, ±3.15mm medial/lateral and -5.00mm dorsal/ventral, relative to bregma (Supplementary Figure S1).

### Tissue oxygen (T_O2_) and Local field potential (LFP) recordings

T_O2_ was measured using constant potential amperometry, as described previously in detail (Lowry *et al*, 2010; McHugh *et al*, 2011; McHugh *et al*, 2013). A constant potential (-650mV relative to a reference electrode) was applied to the electrode, which results in the electrochemical reduction of O_2_ on the electrode surface such that local changes in O_2_ concentration produce directly proportional changes in the measured Faradaic current.

### Fear conditioning

Fear conditioning was conducted in one of two operant chambers (ENV-307A, Med Associates Inc., IN, USA), each with distinct visual and olfactory cues. Mice (n=60; including n=28 who had electrode implantation surgery) underwent a pre-exposure day, three training days (T1, T2, T3), and a fear memory recall day (R).Training was performed in one context (e.g. context A) and recall/extinction was performed in a different context (e.g. context B if trained in context A). Contexts were counterbalanced across mice.

Mice were trained to discriminate between two distinct auditory cues (tone, white noise), with one conditioned stimulus (e.g. CS+ = tone) always paired with foot-shock during the training sessions and the other never paired with shock (e.g. CS- = white noise). Allocation of tone and white noise to the CS+ / CS- was fully counterbalanced across mice. During each session mice were presented with 10 auditory cues (5 × 2900 Hz tone, 5 × white noise; all 72 dB and 30 s duration) in a pseudo-randomly interleaved order with a mean inter-cue interval of 80 s (range 60-100 s). On training days, each of the five CS+ trials co-terminated with mild foot-shock (0.3 mA, 0.5 s). No shocks were given during the pre-exposure or recall sessions.

### Data Analyses

#### Behavior

Freezing responses were assessed by automated movement detection using software running in NIH image (Richmond *et al*, 1998). To analyze CS evoked freezing responses (Δfreezing), we calculated percentage freezing in the 30 s before CS onset and subtracted this from percentage freezing during CS presentation (Figure 1C,D). CS evoked freezing greater than the pre-CS baseline yields positive scores and CS evoked freezing less than pre-CS period yields negative scores (Figure 1C,D). We also calculated a discrimination index by subtracting the CS-Δfreezing score from the CS+ Δfreezing score (Figure 1E). Note that there were no genotypic differences in freezing during pre-CS or CS periods during the pre-exposure session (all F’s < 1.4, all p’s > 0.2; Figure S2). Note also that freezing levels during the pre-CS periods did not differ between genotypes during training or recall (no main effect of genotype or genotype × day interaction: F < 1, p > 0.8). Freezing responses during pre-CS and CS periods for all days are shown in supplementary Figure S3.

**Figure 1.**
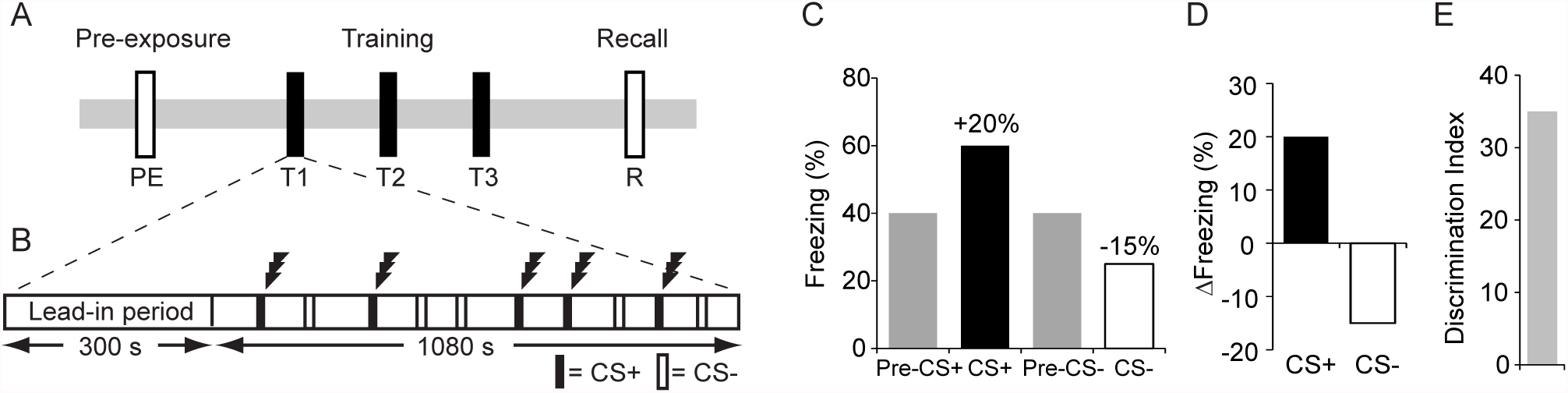
Experimental design. **A** Schematic of the training paradigm. **B** Each session contained a 300s lead-in period followed by five CS+ and five CS- presentations, pseudo-randomly interleaved, and with a different presentation sequence on each day. On training days (T1-T3), the CS+ co-terminated with foot-shock (lightening icon), whereas the CS- was never paired with foot-shock. No shocks were given during pre-exposure (PE) or recall (R) sessions. (**C-D**) Illustration of how percentage freezing scores during pre-CS and CS periods (shown in C) were used to create Δfreezing difference scores [CS minus pre-CS] (shown in D). **E** A discrimination index was generated by subtracting CS- from CS+ Δfreezing scores (e.g. +20 – (-15) = +35).

#### Tissue Oxygen (T_O2_) responses

CS-evoked T_O2_ responses were calculated by subtracting the mean T_O2_ signal during the 5 s before CS onset (i.e. pre-CS baseline) from the T_O2_ signal during the 30 s CS presentation. Then, the 30 s signal was divided into fifteen 2 s time bins (i.e. 0-2 s, 2-4 s, 4-6 s … 28-30 s) with each data point equal to the mean value during each 2 s time bin. The foot-shock (US) was 0.5s duration but we analyzed US-evoked signals in the 30s after the shock, not including the 0.5s in which the US was administered. We averaged T_O2_ responses over the five CS+, CS-, US trials of each session. In addition, we used a regression analysis to investigate whether T_O2_ signals could predict behavioral discrimination, and for this analysis we calculated the maximum T_O2_ signal (i.e. the peak value in one of the 15 time bins) during the CS+, CS- and US periods. The regression model used these values (T_O2__CS+, T_O2__CS-, and T_O2__shock) on training days 1, 2, and 3 to predict behavioral discrimination on subsequent training days 2 and 3 and the recall session, respectively. Stepwise linear regression was used to determine the maximum variance explained with the fewest independent variables.

#### Local field potentials (LFPs)

LFPs were band-pass filtered between 1-80 Hz. We calculated power spectra during the 10s after CS onset for each trial and then averaged these spectra over the five CS+ or five CS- trials in each session. Spectra were computed in MATLAB (The Mathworks, MA, USA). For statistical analysis, the power spectral density in each frequency bin (Φ_i_,) was transformed into a proportion of the total power between 1 and 40 Hz:

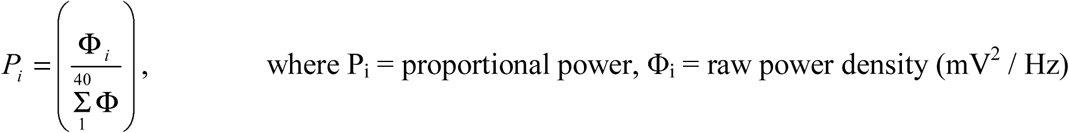

From these proportional spectra, we determined the peak power in the theta frequency range (5-10 Hz).

#### Histology

At the end of the experiments, mice implanted with electrodes were injected with sodium pentobarbitone; 200 mg/kg) and perfused transcardially with physiological saline followed by 10% formol saline. Coronal sections (40 μm) were stained with cresyl violet to determine that electrodes were in the BLA (supplementary Figure S1).

#### Drug treatment

In experiment 2, a separate group of mice (n=96) were injected with saline or citalopram (10mg/kg, i.p.; Tocris, U.K) 30 minutes before each training and/or recall session using a fully counterbalanced design (Table 1). The behavioral paradigm was almost identical to that used in Experiment 1, except there was no pre-exposure session and only two days of training. There were four separate treatment groups: mice given saline during training and recall (Sal_Sal), mice given citalopram during training and saline during recall (Cit_Sal), mice given saline during training and citalopram during recall (Sal_Cit), and mice given citalopram during both training and recall (Cit_Cit).

**Table 1.** Experimental Design for the citalopram study. Mice (n=96) were divided into 4 groups and injected with either saline (Sal) or citalopram (Cit; 10 mg/kg, i.p.) 30 minutes before each training or recall session.

|  | **Training day 1** | **Training day 2** | **Recall** |
| --- | --- | --- | --- |
| **Sal_Sal (n=24)** | Sal | Sal | Sal |
| **Sal_Cit (n=24)** | Sal | Sal | Cit |
| **Cit_Sal (n=24)** | Cit | Cit | Sal |
| **Cit_Cit (n=24)** | Cit | Cit | Cit |

#### Statistical procedures

Data were analyzed using analysis-of-variance (ANOVA) in SPSS (version 22, IBM, USA). ANOVAs are described in the form: A_2_ × B_3_, where A is a factor with two levels and B a factor with three levels. All graphs show the mean ± 1 standard error of the mean (SEM). The familywise error was set at α = 0.05.

## Results

### Facilitated discriminative aversive learning in 5-HTTKO mice

Both genotypes learned to discriminate between the aversive (CS+) and non-aversive (CS-) cues by training day 2 (T2), freezing more during CS+ than CS- trials, but this discrimination was markedly stronger in 5-HTTKO mice (Figure 2A-B). Analysis of Δfreezing scores (ANOVA model: genotype_2_ × sex_2_ × day_2_ × CS type_2_ × trial_5_, n = 60 mice) revealed a main effect of CS type (F(1,56) = 23.4, p<0.001; with CS+ > CS-) and, importantly, superior discrimination in 5-HTTKO versus WTs (genotype × CS type interaction: F(1,56)=7.6, p=0.01). 5-HTTKO mice exhibited significant discrimination during training day 1 (T1) whereas WTs did not (p=0.01 versus p=0.7; see Figure S3). Moreover, analysis of the discrimination index (calculated as [CS+] – [CS-] Δfreezing scores), revealed significantly stronger discrimination in 5-HTTKO mice versus WTs during T2 (F(1,56)=8.0, p=0.006; Figure 2B). Thus, 5-HTTKO mice exhibited facilitated discriminative aversive learning.

**Figure 2.**
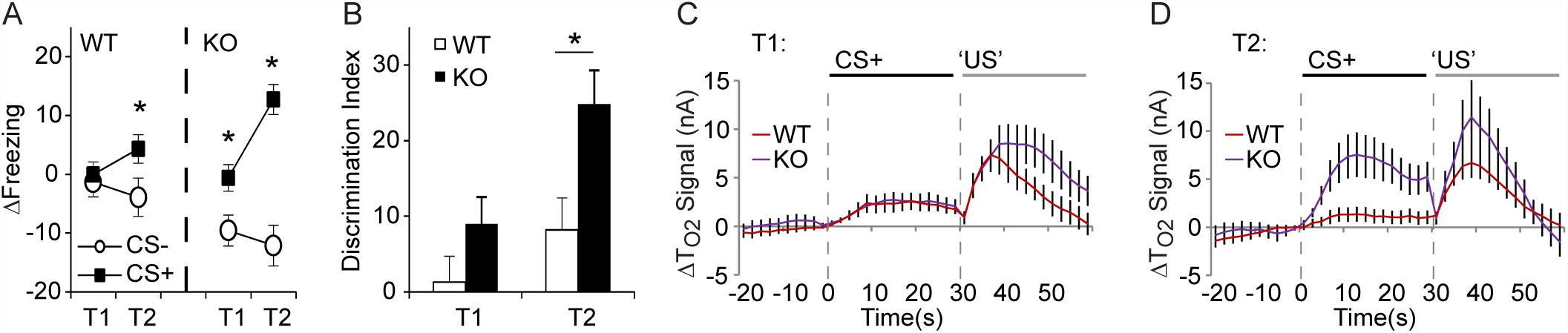
Behavioral (A-B) and amygdala hemodynamic tissue oxygen responses (ΔTO2 signals) (C-D) during discriminative aversive learning in wild-type (WT) and 5-HTTKO (KO) mice. (A) 5-HTTKO mice exhibited larger differences between CS+ and CS- evoked Δfreezing scores during training. Note that positive Δfreezing scores indicate an increase in freezing compared to the pre-CS period whereas negative scores indicate a decrease in freezing compared to the pre-CS period. (B) The discrimination index (CS+ Δfreezing – CS- Δfreezing scores) was significantly higher in 5-HTTKO than WT mice during T2. (C-D) Mean amygdala hemodynamic responses during the 20s before CS onset, 30s during the CS+, and 30s after US onset in WT and 5-HTTKO mice on T1 (C) and T2 (D). Note that TO2 signals were higher in 5-HTTKO mice following foot-shock on T1 and higher during CS+ presentations on T2. (A-B) Mean ± SEM, n=60 mice. (C-D) Mean ± SEM, n=24 mice.*p < 0.05, **p < 0.01.

### Larger cue evoked amygdala hemodynamic responses in 5-HTTKO mice

Tissue oxygen voltammetry measures the hemodynamic response reflecting on-going neuronal activity, similar to the BOLD signal in fMRI. We have shown previously that these hemodynamic responses provide a neural correlate of aversive learning in the amygdala (McHugh et al., 2014). Stimulus-evoked T_O2_ responses in the amygdala were larger in 5-HTTKO than WT mice during fear learning. Analysis of T_O2_ signals during the first two training days (ANOVA: genotype_2_ × stimulus type_3 (CS-, CS+, US)_, × day_2_ × time bin_15_, n=24 mice) revealed that during T1, shock-evoked T_O2_ responses were larger in 5-HTTKO mice (Figure 2C). This higher amygdala signal in 5-HTTKO mice could not be explained by greater locomotor activity in response to the foot-shock (Supplementary Figure S4). During T2, both CS+ evoked and CS- evoked T_O2_ responses were larger in 5-HTTKO mice (Figure 2D; interactions between genotype, stimulus type and day: F(2,44)=3.6, p=0.035, and genotype, stimulus type, day, and time bin: F(28,616)=2.4, p<0.001; for CS- responses, see Supplementary Figure S5). Note that both genotypes had larger T_O2_ responses to the CS+ than the CS-, but this was not apparent in WTs until training day 3 (Supplementary Figure S5).

### Amygdala hemodynamic responses to foot-shock predict subsequent fear-related behavior

Next, we used multiple linear regression to investigate whether T_O2_ signals on day *n* could predict behavioral discrimination between the CS+ and CS- on day *n+ 1* (McHugh *et al*, 2014). The dependent variable was the discrimination index and the three independent variables used were: the maximum T_O2_ signal during CS+ presentations (T_O2__CS+), the maximum T_O2_ signal during CS- presentations (T_O2__CS-), and the maximum T_O2_ signal in the 30s following foot-shock (T_O2__shock). The optimum regression model contained only T_O2__shock and accounted for 7% of the variance in behavioral discrimination, which was significantly greater than zero (F(1,68)=5.1, p=0.03, R^2^=0.07). There was a significant positive correlation between T_O2__shock and behavioral discrimination (standardized β coefficient=0.27; t(68)=2.4, p=0.03). In short, and consistent with a previous report (McHugh *et al*, 2014), mice that exhibited larger T_O2_ responses to foot-shock exhibited better behavioral discrimination the following day. The larger shock-evoked responses in 5-HTTKO than WT mice during T1 (Figure 2C) may therefore explain, at least in part, the superior discrimination in 5-HTTKO mice during T2.

### Larger cue-evoked theta frequency neuronal oscillations in 5-HTTKO mice

Next, we investigated whether the enhanced discrimination learning in 5-HTTKO was accompanied by changes in amygdala oscillatory neuronal activity. Previous work suggests that theta frequency (5-10 Hz) oscillations are an important marker for synaptic plasticity and are evoked by aversive cues (McHugh *et al*, 2014; Seidenbecher *et al*, 2003). Consistent with these previous reports, here we found that during training and recall the onset of the CS+ produced an increase in theta oscillations and a decrease in delta oscillations (1-4 Hz). This increase in theta was evident in both genotypes but was markedly stronger in 5-HTTKOs (Figure 3).

**Figure 3.**
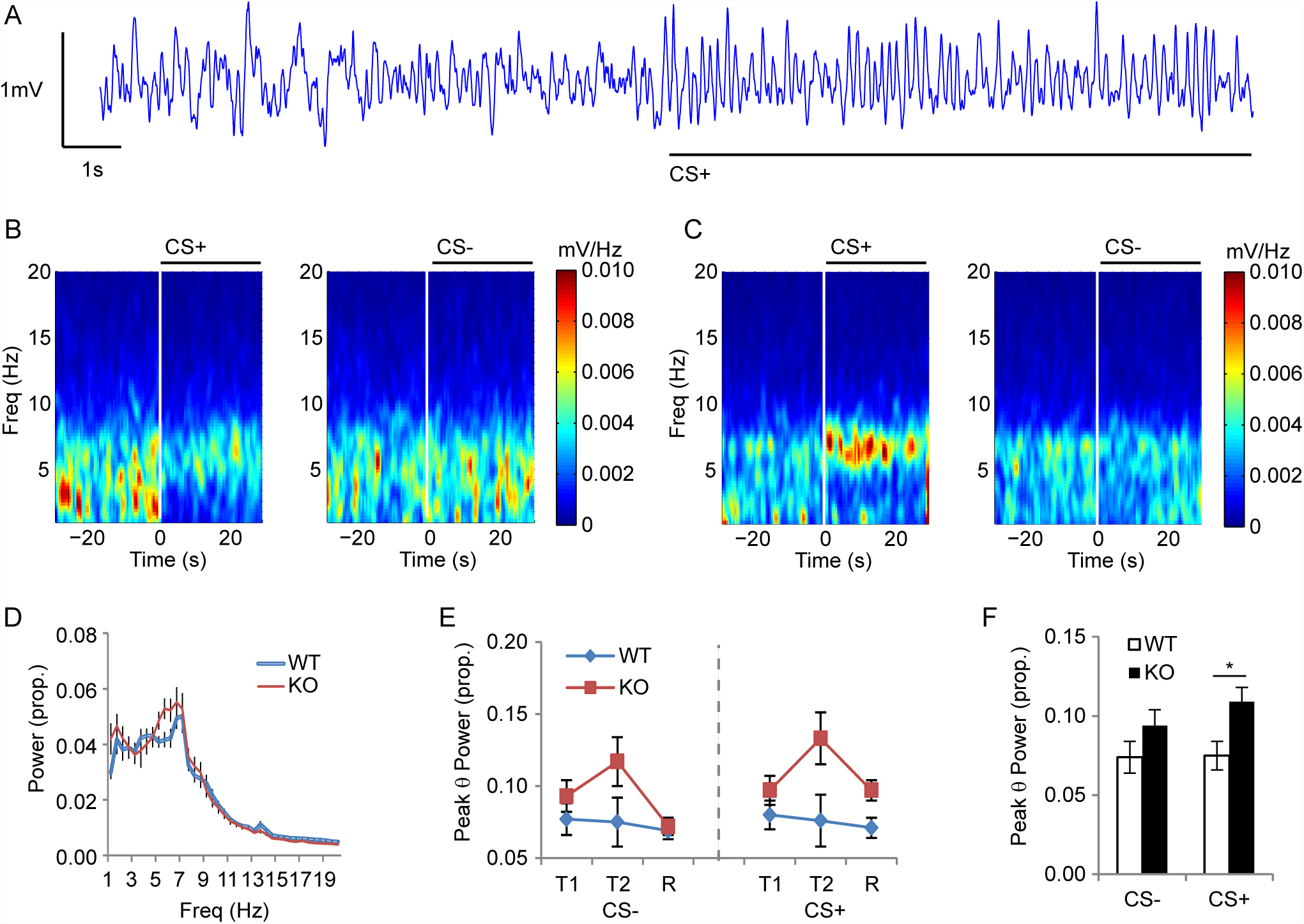
Cue evoked neuronal oscillations in wild-type (WT) and 5-HTTKO (KO) mice during discriminative fear conditioning. (**A**) Raw local field potential trace showing theta activity evoked by CS+ onset. **(B-C)** Time-frequency spectrograms from one WT (B) and one 5-HTTKO (C) mouse during T2 illustrating how CS+ onset evoked a shift from delta-dominant (1-4 Hz) to theta-dominant (5-10 Hz) oscillations. **(D-E)** Peak CS+ evoked theta power was higher in 5- HTTKO than WT mice (D-F) Mean ± SEM, n=24 mice. *p < 0.05.

Analysis of responses during T1, T2 and R (ANOVA: genotype_2_ × day_3_ × CS type_2_, n=24 mice) revealed that theta oscillations were stronger during CS+ than CS- trials (main effect of CS type: F(1,22)=10.8, p=0.003) and were stronger in 5-HTTKO mice than WTs (main effect of genotype: F(1,22)=4.24, p=0.05; genotype × CS type interaction: F(1,22)=7.1, p=0.01). Pairwise comparisons revealed that CS+ evoked theta power was significantly higher in 5-HTTKO mice than WTs (p=0.02), with no genotypic difference for CS- trials (p=0.15). Thus, 5-HTTKO mice exhibited enhanced amygdala theta oscillations during the aversive conditioned cues.

### Citalopram given specifically during training sessions facilitates subsequent fear memory recall

5-HTTKO mice have higher levels of evoked extracellular 5-HT, as measured using microdialysis and fast cyclic voltammetry (Jennings *et al*, 2010; Mathews *et al*, 2004). Given the constitutive nature of the knockout, these mice will experience greater 5-HT availability during development and throughout their entire lifetime. Hence, higher levels of 5-HT availability, either at the time of conditioning or earlier in life, could account for the facilitated fear discrimination in 5-HTTKO mice. To test whether the enhanced discrimination performance reflects elevated 5-HT levels at the time of learning, we trained normal wild-type (C57BL/6J) mice on an almost identical fear conditioning task to that used in the 5-HTTKO mice, with mice receiving the SSRI citalopram (10 mg/kg, i.p.) or saline 30 minutes before each session in a fully counterbalanced design (see Table 1).

Analysis of the discrimination index revealed that citalopram treatment did not affect discrimination during the training days (no effect of day, drug-treatment or interaction, F<1, p>0.5; Figure 4). However, mice that had received citalopram during training exhibited significantly better discrimination during the fear memory recall session, regardless of whether they received saline or citalopram before the fear memory recall session (main effect of treatment during training: F(1,94)=8.5, p=0.005; Figure 4). When comparing performance across the four different treatment groups specifically during fear memory recall (Figure 4B), the Cit_Sal and the Cit_Cit groups exhibited significantly better discrimination than the Sal_Sal group (p=0.001 and p=0.03, respectively); and the Cit_Sal group discriminated better than the Sal_Cit group (p=0.05). There were no other significant group differences. Citalopram did not affect freezing during pre-CS periods (Supplementary Figure S6) or responsivity to the foot-shock, except on the very first trial (Supplementary Figure S7). Thus, mice that received citalopram during training exhibited enhanced fear memory recall. Combined with the data from the 5-HTTKO mice, these results support the hypothesis that increased 5-HT availability at the time of conditioning enhances aversive discrimination learning.

**Figure 4.**
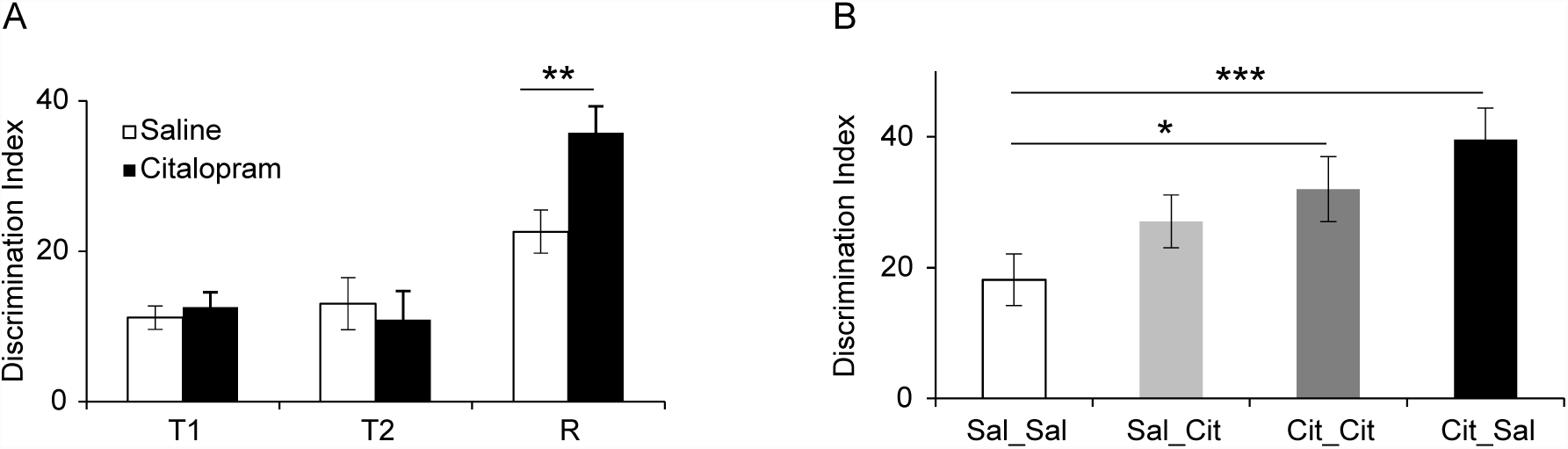
**(A)** Citalopram treatment 30 minutes before training sessions had no effect on CS+ / CS- discrimination during the training sessions (T1, T2) but mice that received citalopram during training exhibited significantly better discrimination during the fear memory recall session (R). Note that during recall, the Saline bar comprises the Sal_Sal and Sal_Cit groups and the Citalopram bar comprises the Cit_Sal and Cit_Cit groups, so the effect was not due to citalopram on fear expression per se (n=96 mice) **(B)** Discrimination during recall was strongest in those mice that received citalopram before training and saline before recall (Cit_Sal) and weakest in mice that received saline before both training and recall (Sal_Sal). ***p=0.001, **p=0.005, *p=0.03.

## Discussion

### Summary of results

We found that 5-HTTKO mice exhibited enhanced discrimination learning for aversive events. Importantly, we also found neural correlates that may underpin these learning effects, with 5- HTTKO mice exhibiting larger amygdala hemodynamic tissue oxygen responses to the aversive cues, including to the foot-shock. Tissue oxygen responses were predictive of subsequent learning, with larger shock-evoked responses predicting better behavioral discrimination between the aversive and non-aversive conditioned cues the following day. Moreover, aversive-cue evoked theta oscillations, a putative marker for increased synaptic plasticity, were stronger in 5- HTTKO mice than WTs. Finally, we found that wild-type mice treated with the SSRI citalopram, specifically before fear training, subsequently exhibited superior fear memory recall. Our data are consistent with network plasticity accounts of 5-HT function, which propose that elevated 5- HT availability increases sensitivity to emotionally-relevant stimuli and therefore enhances learning and memory.

### 5-HTTKO mice have superior aversive learning

Previous studies have reported normal fear learning in 5-HTTKO rodents (Muller *et al*, 2011; Narayanan *et al*, 2011; Shan *et al*, 2014; Wellman *et al*, 2007) but none of these studies used discriminative approaches comparable to the present study. Single-cue fear paradigms typically produce strong fear responses (i.e. high levels of freezing), which can reduce the chances of detecting genotypic differences during fear learning because of ceiling effects. Moreover, discrimination tasks require the mice to attend to the specific sensory features of the auditory cues to a greater extent than single-cue paradigms. In this respect, it is worth noting that the enhanced discrimination performance in the 5-HTTKO mice was driven by responses to both the CS+ and CS-, and was mirrored by larger amygdala hemodynamic responses to both the CS+ and CS-, compared to wild-type controls. Thus, our data show stronger discrimination in 5- HTTKO mice but not increased freezing per se, and are therefore not inconsistent with previous studies.

Findings of improved aversive learning in 5-HTTKO mice complement recent studies showing impaired aversive learning in 5-HTT over-expressing (5-HTTOE) mice (Barkus *et al*, 2014; Line *et al*, 2014; McHugh *et al*, 2015), which have ~3-fold greater 5-HTT expression than their WT counterparts, and lower 5-HT availability (Barkus *et al*, 2014; Jennings *et al*, 2010; Jennings *et al*, 2006). Our results are also consistent with human studies showing superior discriminative aversive learning in putative low 5-HTT expressing genotypes (i.e. SS/SL vs LL) (Garpenstrand *et al*, 2001; Lonsdorf *et al*, 2009) and a positron emission tomography study showing that lower amygdala 5-HTT expression is associated with enhanced discriminative aversive learning (Ahs *et al*, 2015). Collectively, these data argue that aversive learning is influenced across a continuum of 5-HTT expression levels in humans and mice, with superior learning in low- or null-expressing 5-HTT variants. In short, elevated 5-HT availability is associated with enhanced aversive learning.

### 5-HTTKO mice have larger amygdala hemodynamic responses

We found that conditioned aversive cues evoked larger amygdala hemodynamic tissue oxygen responses in 5-HTTKO than WT mice. Our data support human 5-HTTLPR functional neuroimaging studies showing that, compared to LL homozygotes, SS and SL exhibit larger amygdala BOLD responses to emotional faces (Hariri *et al*, 2002). Our tissue oxygen results are also consistent with an *ex vivo* autoradiographic study, which reported higher amygdala blood-flow in 5-HTTKO mice following fear memory recall (Pang *et al*, 2011). Moreover, we have recently shown lower fear-evoked amygdala tissue oxygen responses in 5-HTTOE mice (Barkus *et al*, 2014) and so the present study provides an important evidence that amygdala tissue oxygen responses are bi-directionally modulated by 5-HTT expression levels in mice. Our tissue oxygen data are also important because they indicate a plausible neural substrate (i.e. enhanced amygdala responses to the unconditioned and conditioned stimuli) for the superior aversive learning in 5-HTTKO mice. Moreover, we found that larger tissue oxygen responses to the unconditioned stimulus (foot-shock) were predictive of better discrimination between the CS+ and CS- the following day, arguing that these signals have a meaningful relationship with the observed behavioral output (McHugh *et al*, 2014).

### 5-HTTKO mice have enhanced theta oscillations

We also found that aversive cue-evoked amygdala theta oscillations were enhanced in 5-HTTKO mice. A large body of work argues that theta oscillations are important for synaptic plasticity and learning (Barkus *et al*, 2014; McHugh *et al*, 2014; Seidenbecher *et al*, 2003; Sejnowski and Paulsen, 2006). Our data are consistent with a previous study showing increased fear-evoked theta-frequency synchronization between the amygdala and pre-frontal cortex in 5-HTTKO mice, although this study did not report whether amygdala theta oscillations were larger *per se* (Narayanan *et al*, 2011). Moreover, we have previously reported that fear-evoked amygdala theta oscillations are reduced in 5-HTTOE mice (Barkus *et al*, 2014), which may underlie their impaired aversive learning (Barkus *et al*, 2014; Line *et al*, 2014; McHugh *et al*, 2015). Theta oscillations are thought to be modulated by the activity of parvalbumin-expressing inhibitory interneurons (PV INs) (Amilhon *et al*, 2015; Wulff *et al*, 2009) and amygdala PV INs are required for successful fear conditioning (Wolff *et al*, 2014). Recently we have shown that both fear conditioning and local 5-HT application activate amygdala PV INs, and that this activation is reduced in 5-HTTOE mice (Bocchio *et al*, 2015). Thus, PV+ INs may be a crucial target by which 5-HT modulates theta oscillations and hence influences fear learning. The enhanced theta in the 5-HTTKO mice is consistent with enhanced recruitment of PV+ INs in these animals.

### Acute SSRI treatment enhances fear learning

Our finding that the SSRI citalopram facilitates fear memory recall is consistent with previous rodent studies (Burghardt *et al*, 2004; Ravinder *et al*, 2013). However, our data extend previous findings by showing, specifically, that citalopram enhances discrimination between the CS+ and CS- and does not simply increase freezing to a conditioned aversive cue. Thus, is the effect of citalopram on recall is unlikely to be due to increased anxiety evoked by acute SSRI administration during training, because anxiety typically increases stimulus generalization (Kheirbek *et al*, 2012; Lissek *et al*, 2010), and might therefore be expected to impair, rather than enhance, discrimination between the CS+ and CS-. In contrast, our data show that elevated 5-HT availability at the time of learning enhances subsequent CS+ / CS- discrimination during recall.

### Conclusions

For decades, the role of 5-HT (and the 5-HTT) in anxiety and depression has proved elusive. The paradox that increased 5-HT levels in experimental animals are anxiogenic and yet SSRIs are effective anxiolytics has proved difficult to resolve (Deakin and Graeff, 1991). The network plasticity theory proposes that, rather than controlling emotion directly, 5-HT’s main role is to facilitate plasticity in the brain circuits that underlie emotional learning (Branchi, 2011; Castren, 2005). This theory is attractive because it can potentially reconcile seemingly contradictory findings. However, until now few studies have provided direct evidence that altering 5-HT availability affects plasticity and/or emotional learning. The present study provides novel behavioral and neural evidence that increasing 5-HT availability enhances learning for emotionally-relevant cues. The relationship between 5-HTT expression levels and the risk of developing affective disorders continues to generate interest, but remains controversial despite two decades of studies. By identifying the psychological and neurobiological mechanisms that are affected by changes in 5-HTT activity, we may be better placed to understand the contribution of 5-HTT activity to anxiety and depression.

### Funding and disclosure

This study was supported by the Wellcome Trust (Grant No. 087736 to D.M.B.), FCT Portugal (doctoral grant SFRH/BD/87496/2012 to J.L.), and Merton College, Oxford (Graduate Research Grants to J.L.).

